## Supplementary File 2 for "The evolution and biological correlates of hand preferences in anthropoid primates"

### Supplementary File 1: Phylogenetic Bayesian regression models on individual lateralization parameters

**Abbreviations:**

Est. Error: Estimate error

l-95% CI: lower limit of 95% credible interval

u-95% CI: upper limit of 95% credible interval

Rhat: Potential scale reduction factor

Bulk_ESS: bulk effective sample size

Tail_ESS: tail effective sample size

Notable effects were assumed when the model's 95% credible intervals of intercept and respective regression coefficients did not overlap with zero (marked in bold).

Figures visualize estimates for individual predictors on the left, and the number of chains and performed iterations on the right.

**Model 1** (Lateralization direction, total sample)

| Group-level effects | | | | | | | |
|---|---|---|---|---|---|---|---|
| | Estimate | Est. Error | l-95% CI | u-95% CI | Rhat | Bulk_ESS | Tail_ESS |
| sd(Intercept) | 0.02 | 0.01 | 0.01 | 0.04 | 1.00 | 1376 | 2057 |
| Population-level effects | | | | | | | |
| Intercept | -0.02 | 0.10 | -0.22 | 0.17 | 1.00 | 1530 | 1565 |
| Sex:male | 0.00 | 0.03 | -0.06 | 0.06 | 1.00 | 5086 | 2613 |
| Age:subadult | -0.07 | 0.04 | -0.15 | 0.00 | 1.00 | 5838 | 3386 |

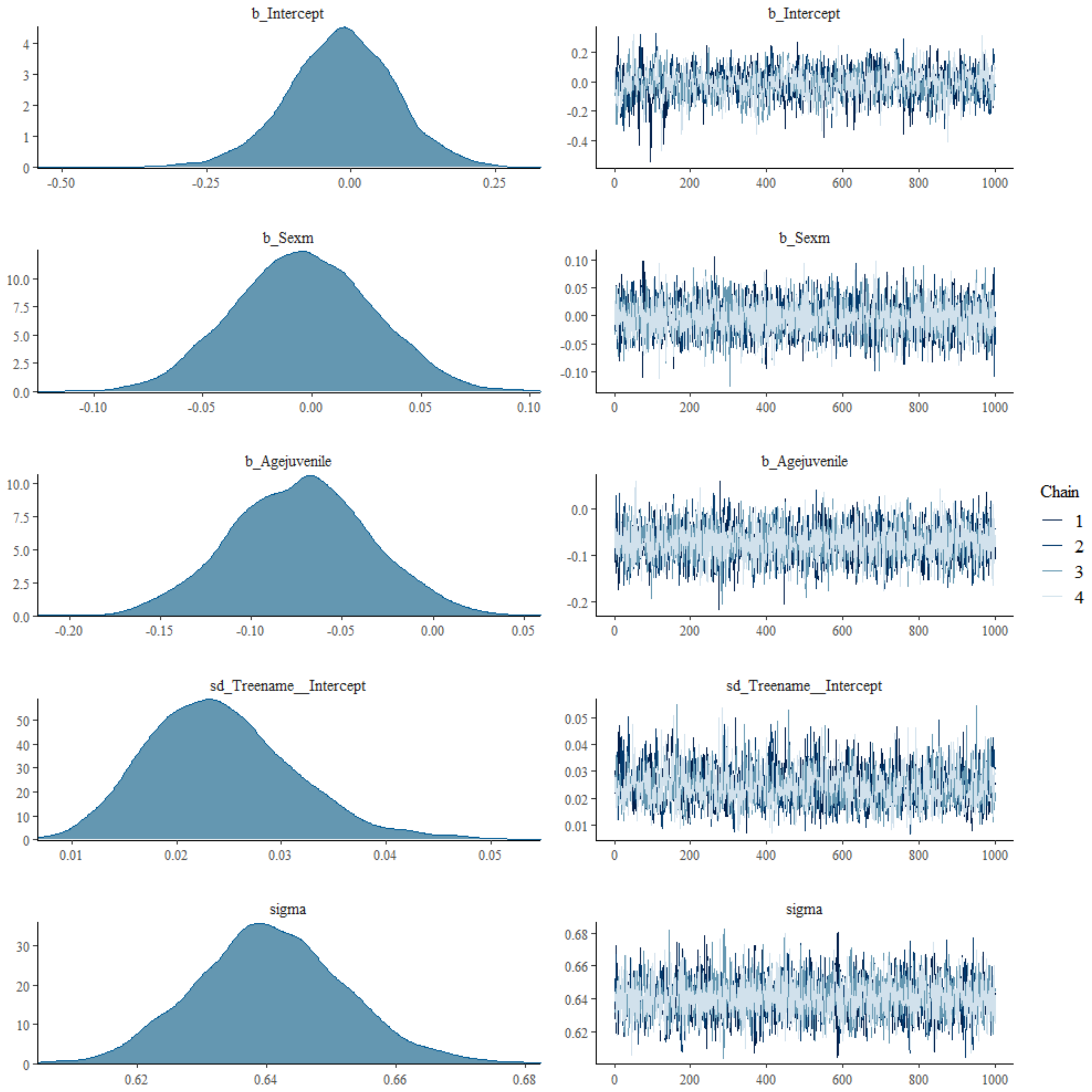

**Model 2** (Lateralization strength, total sample)

| Group-level effects | | | | | | | |
|---|---|---|---|---|---|---|---|
| | Estimate | Est. Error | l-95% CI | u-95% CI | Rhat | Bulk_ESS | Tail_ESS |
| sd(Intercept) | 0.04 | 0.01 | 0.03 | 0.05 | 1.00 | 1149 | 1835 |
| Population-level effects | | | | | | | |
| Intercept | 0.67 | 0.14 | 0.40 | 0.93 | 1.00 | 829 | 1754 |
| Sex:male | 0.02 | 0.02 | -0.01 | 0.05 | 1.00 | 4560 | 3068 |
| **Age:subadult** | **-0.08** | **0.02** | **-0.11** | **-0.05** | **1.00** | **4581** | **3076** |

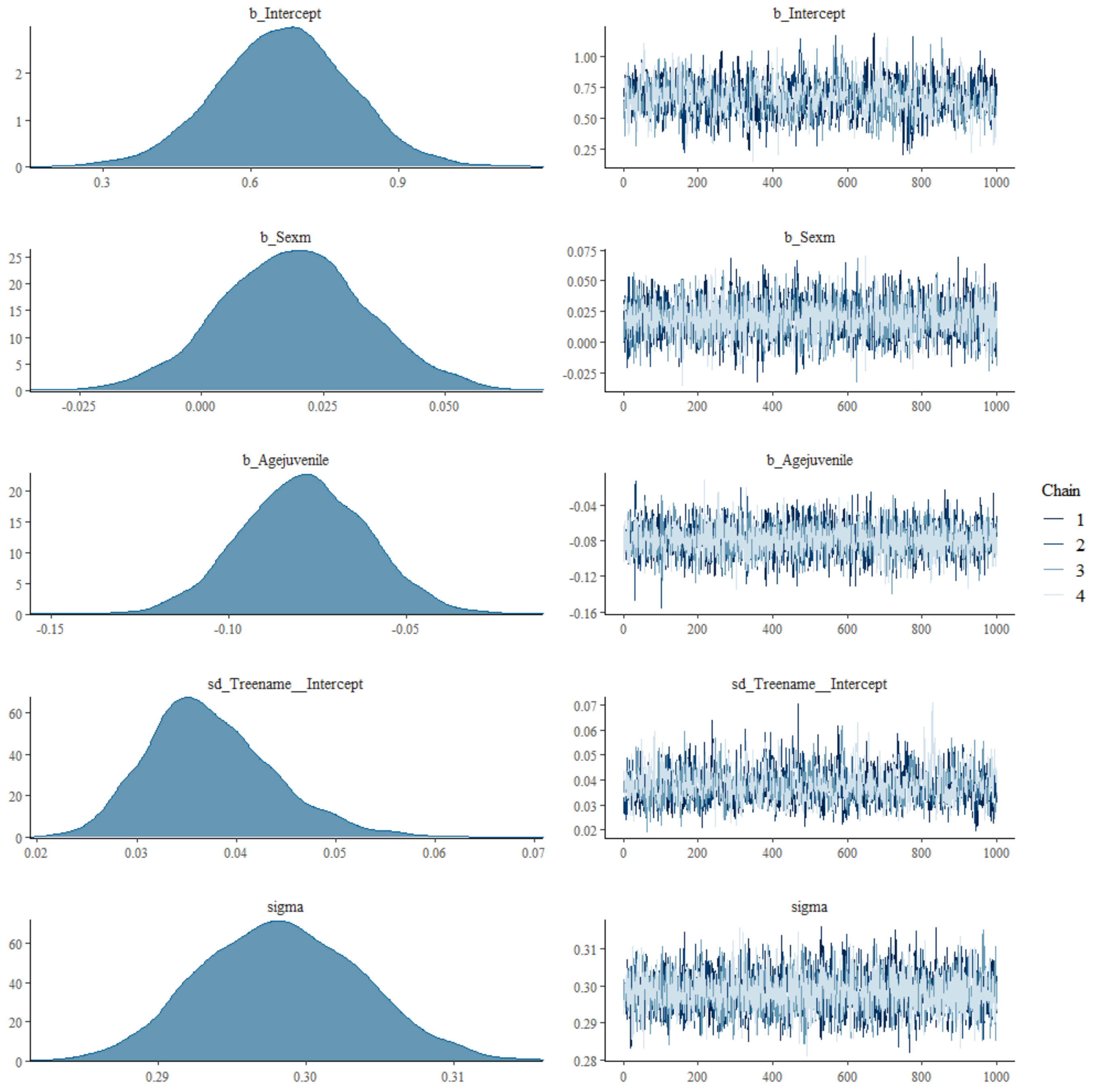

**Model 3** (Lateralization direction, Platyrrhini)

| Group-level effects | | | | | | | |
|---|---|---|---|---|---|---|---|
| | Estimate | Est. Error | l-95% CI | u-95% CI | Rhat | Bulk_ESS | Tail_ESS |
| sd(Intercept) | 0.02 | 0.02 | 0 | 0.08 | 1.00 | 1717 | 784 |
| Population-level effects | | | | | | | |
| Intercept | 0.05 | 0.16 | -0.28 | 0.38 | 1.01 | 1005 | 615 |
| Sex:male | -0.11 | 0.09 | -0.30 | 0.07 | 1.00 | 6417 | 6563 |
| Age:subadult | -0.13 | 0.14 | -0.41 | 0.15 | 1.00 | 9115 | 7120 |

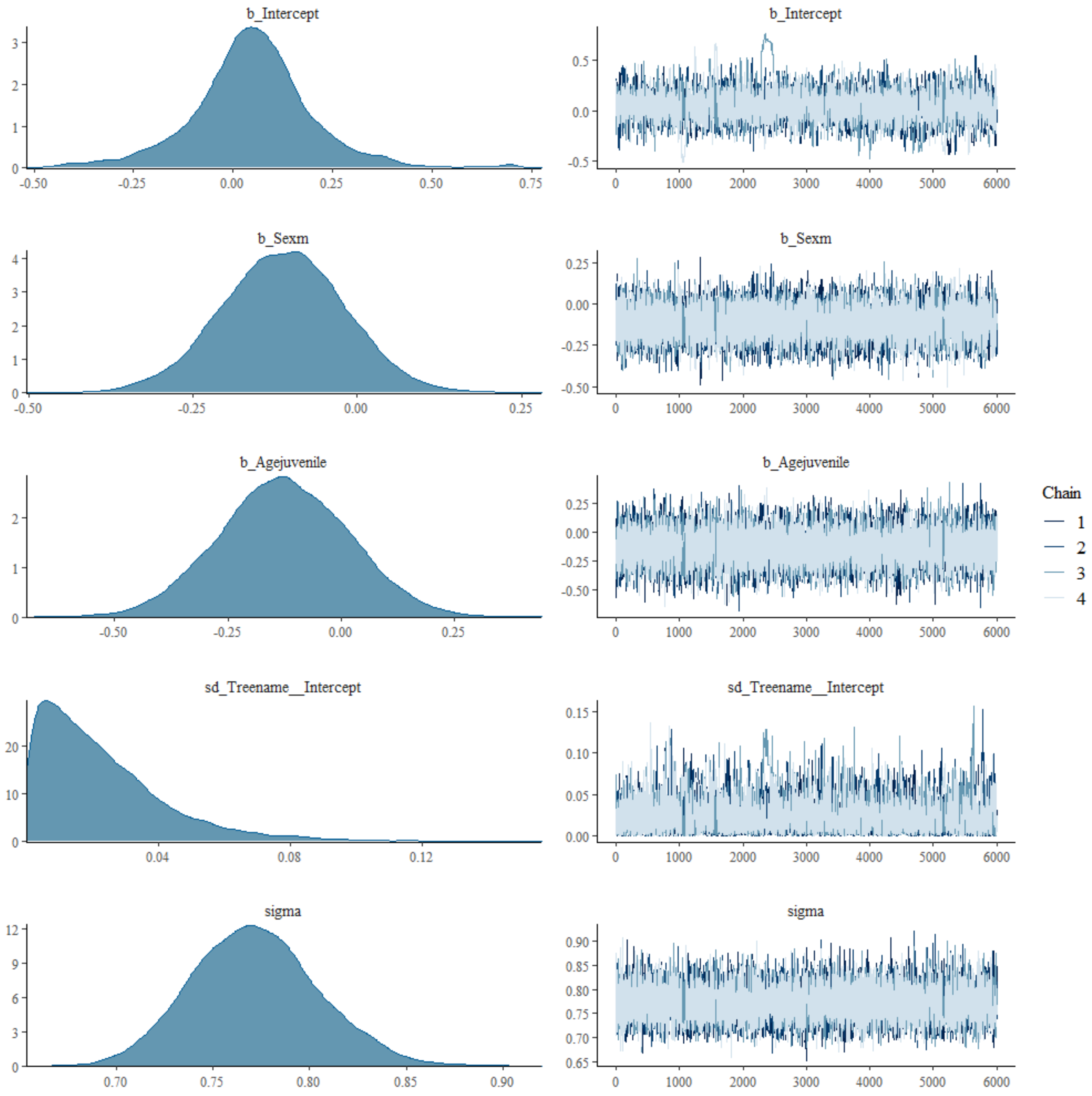

**Model 4** (Lateralization strength, Platyrrhini)

| Group-level effects | | | | | | | |
|---|---|---|---|---|---|---|---|
| | Estimate | Est. Error | l-95% CI | u-95% CI | Rhat | Bulk_ESS | Tail_ESS |
| sd(Intercept) | 0.04 | 0.02 | 0.02 | 0.08 | 1.00 | 1692 | 1048 |
| Population-level effects | | | | | | | |
| Intercept | 0.76 | 0.23 | 0.29 | 1.21 | 1.00 | 1854 | 877 |
| Sex:male | -0.05 | 0.04 | -0.12 | 0.02 | 1.00 | 13737 | 6729 |
| Age:subadult | -0.03 | 0.05 | -0.14 | 0.07 | 1.00 | 7848 | 2185 |

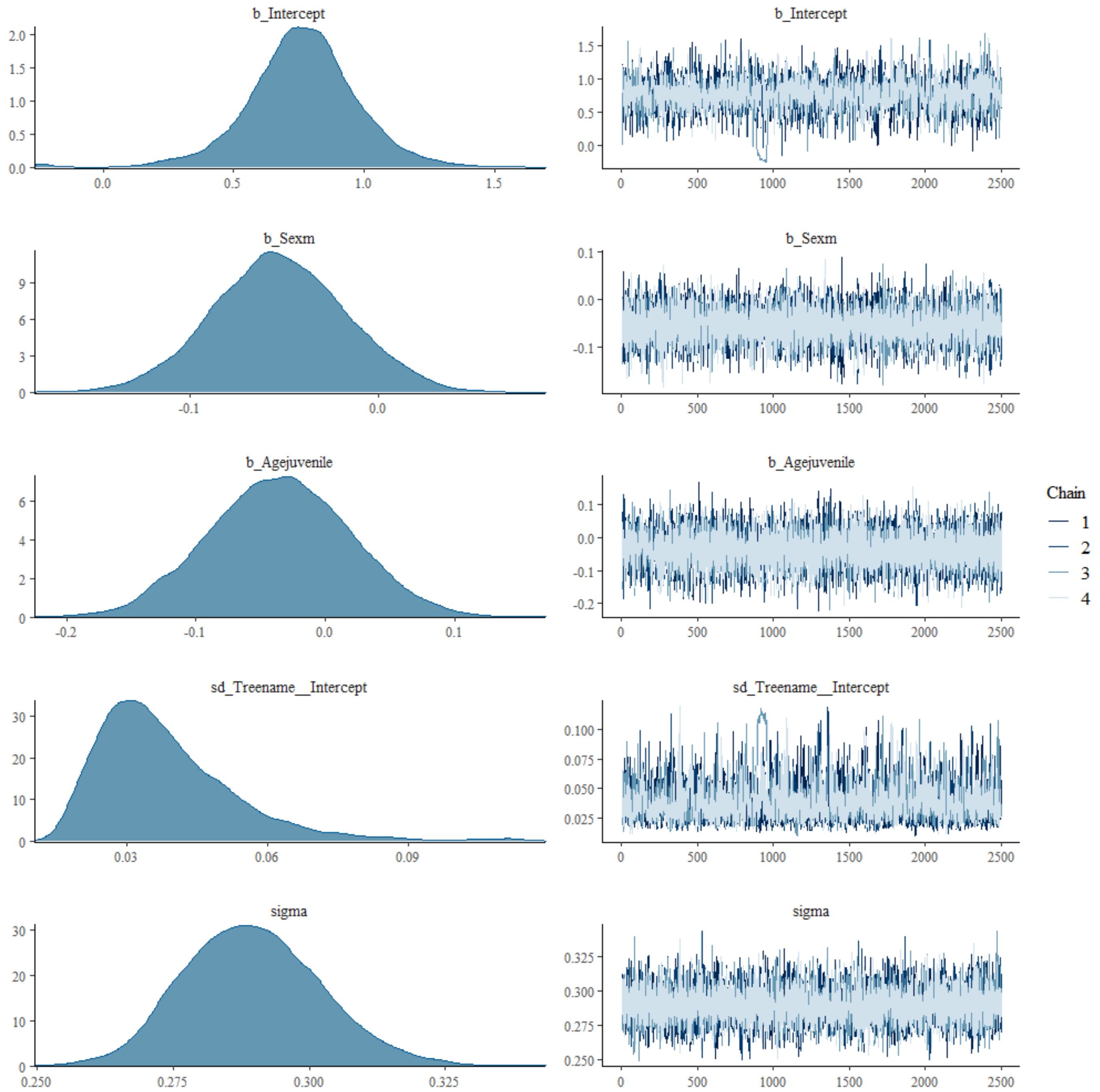

**Model 5** (Lateralization direction, Cercopithecoidea)

| Group-level effects | | | | | | | |
|---|---|---|---|---|---|---|---|
| | Estimate | Est. Error | l-95% CI | u-95% CI | Rhat | Bulk_ESS | Tail_ESS |
| sd(Intercept) | 0.02 | 0.01 | 0.09 | 0.05 | 1.01 | 640 | 419 |
| Population-level effects | | | | | | | |
| Intercept | -0.06 | 0.17 | -0.48 | 0.27 | 1.01 | 541 | 194 |
| Sex:male | 0.02 | 0.06 | -0.10 | 0.14 | 1.00 | 3062 | 1473 |
| Age:subadult | -0.08 | 0.09 | -0.25 | 0.09 | 1.00 | 3388 | 2618 |

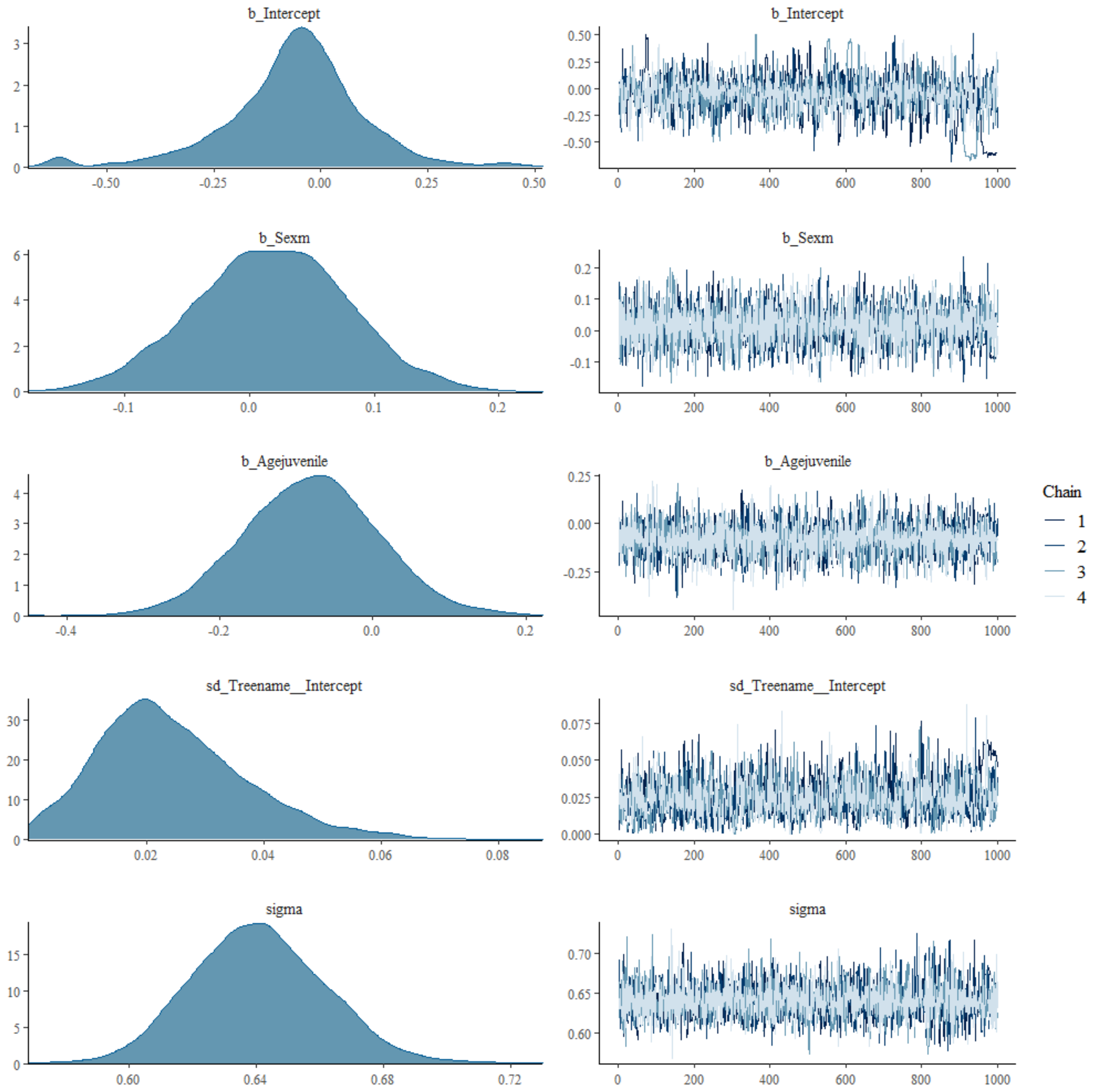

**Model 6** (Lateralization strength, Cercopithecoidea)

| Group-level effects | | | | | | | |
|---|---|---|---|---|---|---|---|
| | Estimate | Est. Error | l-95% CI | u-95% CI | Rhat | Bulk_ESS | Tail_ESS |
| sd(Intercept) | 0.05 | 0.01 | 0.03 | 0.08 | 1.01 | 772 | 1284 |
| Population-level effects | | | | | | | |
| Intercept | 0.60 | 0.30 | 0 | 1.20 | 1.00 | 1060 | 1406 |
| Sex:male | 0.05 | 0.03 | 0 | 0.11 | 1.00 | 3981 | 2986 |
| Age:subadult | -0.04 | 0.04 | -0.12 | 0.04 | 1.00 | 4070 | 3171 |

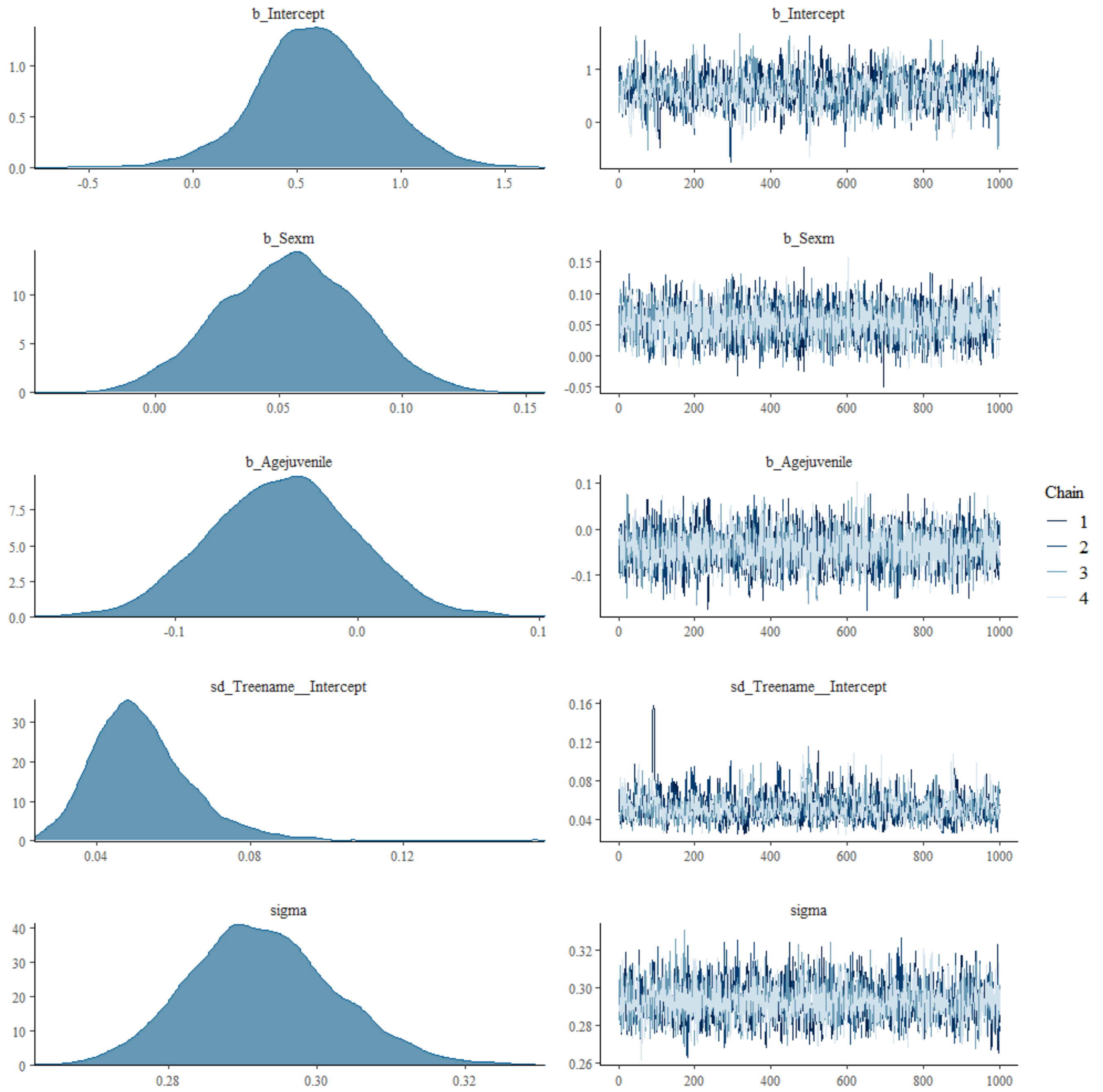

**Model 7** (Lateralization direction, Hominoidea)

| Group-level effects | | | | | | | |
|---|---|---|---|---|---|---|---|
| | Estimate | Est. Error | l-95% CI | u-95% CI | Rhat | Bulk_ESS | Tail_ESS |
| sd(Intercept) | 0.05 | 0.02 | 0.02 | 0.10 | 1.00 | 1862 | 908 |
| Population-level effects | | | | | | | |
| Intercept | 0.04 | 0.30 | -0.55 | 0.70 | 1.00 | 1806 | 850 |
| Sex:male | 0.04 | 0.07 | -0.09 | 0.17 | 1.00 | 9879 | 7468 |
| **Age:subadult** | **-0.15** | **0.07** | **-0.28** | **-0.01** | **1.00** | **8675** | **7153** |

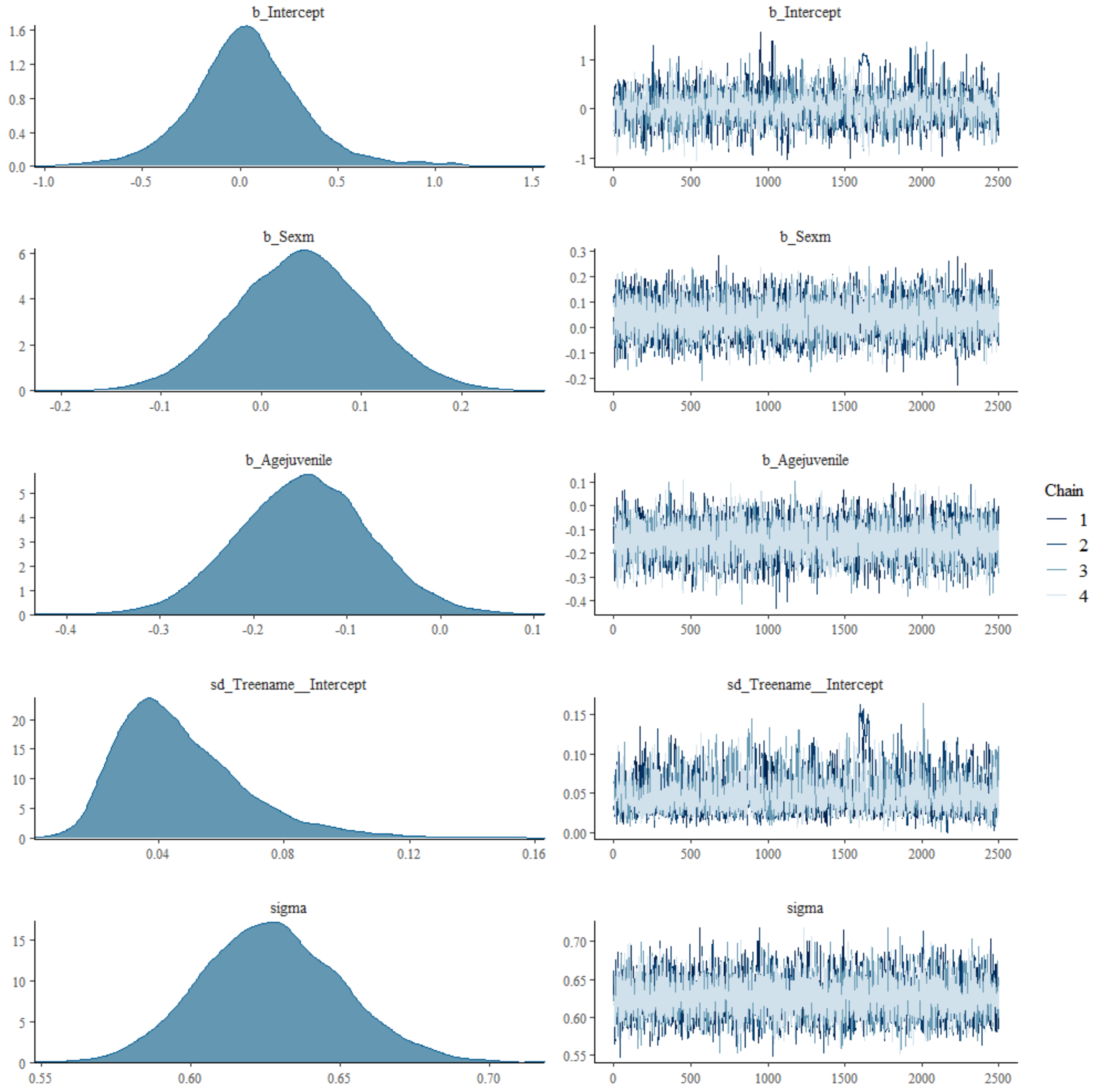

**Model 8** (Lateralization strength, Hominoidea)

| Group-level effects | | | | | | | |
|---|---|---|---|---|---|---|---|
| | Estimate | Est. Error | l-95% CI | u-95% CI | Rhat | Bulk_ESS | Tail_ESS |
| sd(Intercept) | 0.03 | 0.02 | 0.00 | 0.07 | 1.00 | 1082 | 903 |
| Population-level effects | | | | | | | |
| Intercept | 0.56 | 0.16 | 0.17 | 0.88 | 1.01 | 754 | 240 |
| Sex:male | 0.03 | 0.03 | -0.03 | 0.10 | 1.00 | 5377 | 8507 |
| **Age:subadult** | **-0.09** | **0.04** | **-0.16** | **-0.02** | **1.00** | **3684** | **5352** |

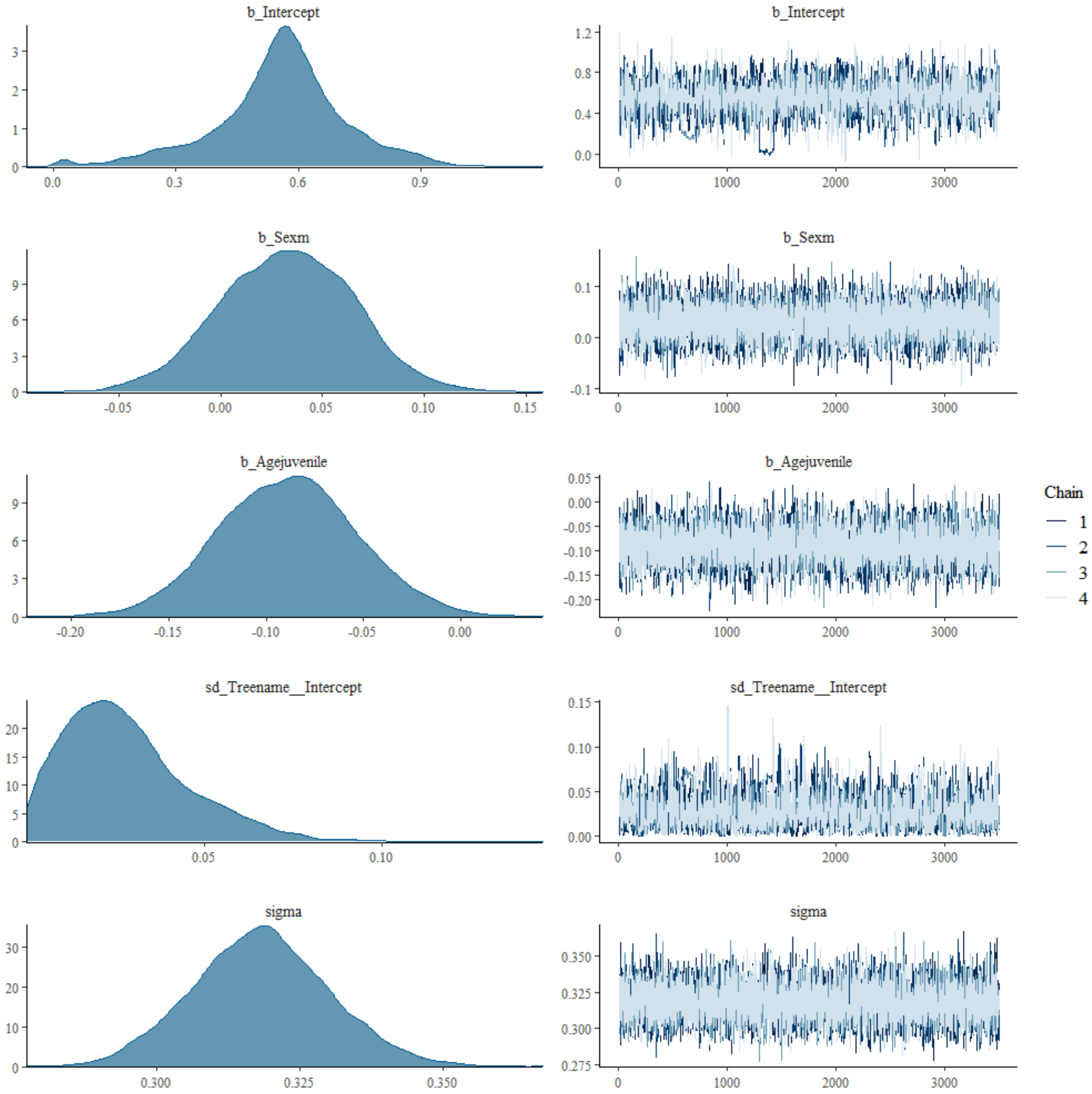
